# A Deep Learning Model for Molecular Label Transfer that Enables Cancer Cell Identification from Histopathology Images

**DOI:** 10.1101/2021.03.18.436004

**Authors:** Andrew Su, HoJoon Lee, Xiao Tan, Carlos J. Suarez, Noemi Andor, Quan Nguyen, Hanlee P. Ji

**Affiliations:** Institute for Molecular Bioscience, The University of Queensland, QLD 4072, Australia; Division of Oncology, Department of Medicine, Stanford University School of Medicine, Stanford, CA, 94305, United States; Department of Pathology, Stanford University School of Medicine, Stanford, CA, 94305, United States; Stanford Genome Technology Center, Stanford University, Palo Alto, CA, 94304, United States

**Author notes:** These authors contributed equally to this work (Andrew Su, HoJoon Lee, Xiao Tan). Current address: Department of Integrated Mathematical Oncology, Moffitt Cancer Center, 12902 Magnolia Drive, Tampa, FL 33612, USA. To whom correspondence should be addressed. Quan Nguyen, Division of Genetics and Genomics, Institute for Molecular Bioscience, The University of Queensland, QLD 4072, Australia, Hanlee P. Ji, Division of Oncology, Department of Medicine – Stanford University School of Medicine CCSR 1115, 269 Campus Drive, Stanford, CA 94305-5151.

**Keywords:** Colorectal cancer, digital pathology, deep learning, cancer diagnosis, computer-assisted tools, histopathological images, molecular label, label transferring

## Abstract

Deep learning cancer classification systems have the potential to improve cancer diagnosis. However, development of these computational approaches depends on prior annotation through a pathologist. This initial step relying on a manual, low-resolution, time-consuming process is highly variable and subject to observer variance. To address this issue, we developed a novel method, H&E Molecular neural network (HEMnet). This two-step process utilises immunohistochemistry as an initial molecular label for cancer cells on a H&E image and then we train a cancer classifier on the overlapping clinical histopathological images. Using this molecular transfer method, we show that HEMnet accurately distinguishes colorectal cancer from normal tissue at high resolution without the need for an initial manual histopathologic evaluation. Our validation study using histopathology images from TCGA samples accurately estimates tumour purity. Overall, our method provides a path towards a fully automated delineation of any type of tumor so long as there is a cancer-oriented molecular stain available for subsequent learning. Software, tutorials and interactive tools are available at: https://github.com/BiomedicalMachineLearning/HEMnet

## BACKGROUND

Histopathological examination of tissue is indispensable for the accurate diagnosis and treatment of cancer^1–3^. Frequently, pathologic diagnosis of cancer and different subtypes dictate the use of specific treatment regimens^4^. One of the current standards of cancer diagnosis is microscopic examination of tumour tissue sections jointly stained with hematoxylin and eosin **(H&E)** dyes^2,3^. Based on the H&E stained image of a biopsy section, pathologists can qualitatively assess cancer types, stages and estimates of tumor purity^3^. Furthermore, histopathologic examination frequently reports different types of cells, organic states, and/or cellular localization inside complex tissues^5^.

The visual inspection of histopathologic sections of biopsies remains a time-consuming task with a high degree of observer variability among pathologists, batch effects from the staining procedures and a lack of quantitative measurements for cellular features^4^. Recently, the emerging area of digital pathology has been developed as a way to digitize, store and distribute cancer whole slide images **(WSIs)**. This approach significantly improves the speed and access to cancer anatomical pathology. The increasing production of WSIs requires advanced computational approaches to be developed to analyze these medical images in a fast, robust and accurate manner, ultimately leading to applications in automated cancer diagnosis^6–9^.

Deep learning is the method of choice for analysis of histology images and has been recently applied to tumour classification on histopathology images^7^. A key challenge for deep learning is the need for large amounts of accurately labeled data. For this approach, many methods require WSIs which are manually annotated by a pathologist. Thus, generating the training data set becomes a time-consuming manual process. This adds to the cost and makes it more expensive to obtain these datasets. Another challenge is that these slide images are large; an image at 10x magnification can contain hundreds of millions of pixels. However, a pathologist’s annotations are often not at the pixel level and rely on much cruder methods of demarcation. As a result, training occurs at a lower image resolution that lacks cellular granularity.

Herein, we describe a new approach in which we use prior staining that demarcates tumors from normal cells at much higher image resolution. For this proof-of-concept study, we used an immunohistochemistry **(IHC)** marker for cancer to delineate tumor cells. Referred to as H&E molecular neural network **(HEMnet)**, this approach increased the size of training dataset at cellular level. The coupling of H&E and molecular marker staining images is increasingly being applied for histopathological evaluation, creating a valuable opportunity for data integration^2,10,11^. In this study, we used p53 staining, an important tumor suppressor gene (*TP53*) which is prone to a high frequency of genetic alterations across many different malignancies^12,13^. Most *TP53* mutations are of the missense class that change the p53 protein structure and lead to their retention in the malignant cell’s cytoplasm. This results in the stabilization and subsequently accumulation of p53, allowing it to be readily detected by IHC. In normal cells, the level of wild-type p53 is usually present in low concentrations undetectable by IHC^14^. In contrast, up to 74% of colorectal cancer samples show abnormal positive staining (i.e. a brown color) for p53, which provides specific IHC marker for cancer cells in colorectal cancer^13,15,16^.

Our study leveraged innovative molecular label transferring to generate tens of thousands of H&E tiles extracted from the WSIs. So long as the molecular label is relatively specific to the tumor cells, this process enables one to conduct streamlined molecular annotation of cancer versus normal cells without the manual inspection. With thousands of labelled tiles, a convolutional neural network classifier was trained based on an in-house colorectal cancer dataset and was tested using public data from the Cancer Image Archive **(TCIA)** database, where H&E images of cancer tissues are accessible^17,18^. We used aberrant TP53 staining patterns to annotate cancer cells in H&E slides by aligning these images. HEMnet was trained on a set of p53-stained and H&E WSI images from colon cancer. With this training approach, we achieved a high performance on an independent set of histopathologic sections and images. HEMnet was extended to testing TCIA colorectal cancer imaging data and by comparing with other genomics-based methods, we demonstrated a high performance with a significant positively correlation. The HEMnet approach can be easily implemented with other interesting biomarkers such as HER2 and for other types of cancer. These developments by multiple molecular markers would enable the analysis of the complexity of the cancer to a greater extent. Given its success this method has potential clinical application through the discovery of cancer cellular geometric patterns within the tissue and the production of software capable of automatic detection of these patterns as part of developing computer aided diagnosis tool.

## MATERIALS AND METHODS

### H&E and IHC image dataset generation

We collected cancer tissue samples from 30 patients at Stanford Hospital. All patients were enrolled according to a study protocol approved by the Stanford University School of Medicine Institutional Review Board (IRB-11886). Informed consent was obtained from all patients. Tissues were obtained from the Stanford Cancer Institute Tissue Bank. In addition, we obtained matched normal, non-cancer tissue from 5 of these patients. Each sample was formalin fixed and paraffin embedded **(FFPE)** as a tissue block and two adjacent sections were taken from each block, ensuring these sections would close to identical. One section was prepared with H&E staining and the other with immunohistochemistry **(IHC)** staining against p53. All digital slide images were generated in Aperio SVS format by Translational Pathology Core Laboratory at University of California, Los Angeles. This study was conducted in compliance with the Helsinki Declaration.

Each tissue section was scanned at 20x magnification to generate a total of 35 p53 and H&E pairs of high resolution WSIs. Pathology review provided the cancer versus normal cell status of these tissues. Three samples stained positive for p53 despite no histopathologic indications of tumor cells, which would have led to inaccurate labelling and model misclassification. To ensure accurate model training and testing, the p53 and H&E WSIs from these samples were excluded in the analysis. Overall, this left a total of 32 pairs of H&E and p53 WSIs, 27 cancer and five normal tissues.

### Training, validating and testing dataset generation

We use a common practice in machine learning of splitting our dataset of WSIs into training, validation and test sets. No overlap existed between these datasets to ensure that test and validation data was completely independent. We assigned the five normal WSI pairs and five cancer WSI pairs to the training dataset. To ensure an accurate training data set, we also confirmed that most p53 stained regions were cancer in these slides by a pathologist. Together, this provided the model the optimal degree of learning to distinguish between cancer and non-cancer tissue **(Supplementary Figure S1a)**. The WSIs were captured at gigapixel scale **(Supplementary Figure S1b)** allowing us to employ a tiling strategy to split each WSI into thousands of smaller 224px x 224px image tiles for neural network training. We set aside five cancer WSI pairs as a validation dataset to optimize our model’s hyperparameters. The remaining 17 cancer WSIs were assigned to an independent test dataset to assess our model’s performance on unseen slides.

### H&E stain colour normalization

Undesirable colour variations occur in H&E staining and imaging due to different immunohistochemistry reagents, protocols and slide scanners ^19^. Therefore, the same cellular structures in a tissue can appear different depending on how the tissue was stained and imaged. To ensure our model generalized to images from H&E slides across different facilities, we corrected for technical variations in the staining and imaging process. First, we corrected for imaging brightness and ensured that the slide background is white through luminosity standardization **(Supplementary Figure S2)**. Next, we normalized each H&E WSI to a reference stain colour profile derived from a template WSI using the Vahadane, et al. ^19^ stain normalization method implemented in StainTools ^20^, described below:

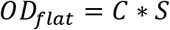

The *OD*_*flat*_ is the flattened optical density **(OD)** array derived from the RGB WSI. A stain matrix (*S*) encodes the stain colour for the H&E staining and is estimated using the Vahadane method. This stain matrix is used to find the pixel stain concentration matrix (*C*). To normalize a source WSI to a template WSI, the stain and concentration matrix for both images are calculated:

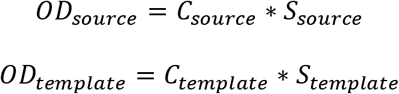

The *C*_*source*_ matrix describes the concentration of hematoxylin and eosin stain at each pixel. Using the stain matrix from the template image (*S*_*template*_) we coloured each pixel in source concentration matrix to produce an image, as if the source image was stained and captured the same way as the template image:

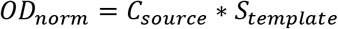

By normalizing all WSIs, training and unseen, to the template image, we ensured that similar cellular structures have the similar appearances regardless of how they were stained and underwent image scanning.

To select a suitable template WSI, we find the cancer slide with mean R, G, B channel intensities closest to the median of the mean of the different channel (R, G and B) intensities of all images **(Supplementary Figure S1c)**. In addition, we implemented two user-selectable, popular but less advanced, image normalization methods by Reinhard, et al. ^21^ and Macenko, et al. ^22^.

### Registration of IHC images to H&E images

For the IHC images to be used to accurately label the H&E images, each IHC image was aligned with its corresponding H&E image. Despite originating from adjacent sections of the same tissue block, technical differences in sectioning, mounting and imaging caused misalignment between IHC images and their H&E counterparts. We aligned these images by implementing image registration through the SimpleITK package^23^.

During registration, the IHC images were warped such that they were aligned to the H&E images. By only transforming the IHC images we ensured that the H&E images remained unaltered. Technical variation among H&E images, for example the variation in the brightness, or color intensities due to microscopy exposure time and/or staining time, was normalised **(Figure S2 and Figure 2)**. Thus, a neural network trained on these H&E images can be applied to new normalised, but otherwise unmodified, H&E images.

We verified the accurate registration through visual inspection and a quantitative mutual information metric. We overlaid the registered p53 over the corresponding H&E image to visually check for correct alignment. In addition, we compared the alignment of p53 image to the H&E image by computing the mutual information between these images before, during and after registration. Mutual information is an information theory concept that can be applied to measure image registration performance **(Supplementary Figure S3)**. An increase in mutual information after registration is indicative better image alignment. The mutual information between the IHC and H&E image can be calculated by:

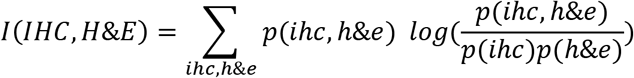

Where p(ihc) and the p(h&e) are the marginal probability distributions of grayscale pixel intensities in the IHC and H&E image respectively. The p(ihc, h&e) is the joint distribution of the images’ grayscale pixel intensities.

Registration strategies can broadly be segregated into feature-based and intensity-based methods. Feature-based methods extract features (e.g. corners) or fiducials from the source and target image and transform the source image such that features in the source image are in the same location as matching features in the target image. On the other hand, intensity-based methods consider the pixel intensity or intensity distributions. These methods also transform the source image such that it most closely correlates with the pixel intensities or intensity distributions of the target image, as measured by a cost function. In preliminary testing, we found that an intensity-based approach was effective for H&E images.

For our intensity-based registration approach, we selected a mutual information cost function to quantify the extent of registering the source and target images. This cost function measures the mutual information between the pixel intensity distributions of the source and target image. The goal of registration is to transform the source image such that the mutual information between the source and target image is maximised - this would imply a well registered image. The mutual information is calculated from grayscale pixel intensities so the IHC and H&E stained images were first converted to grayscale. Post-registration, the optimal transform for the grayscale IHC image is applied to each channel of the RGB IHC image to produce a registered colour image.

To achieve accurate registration and reach a global, rather than local, optima, we performed affine registration followed by b-spline registration. The initial linear affine registration is limited to translation, scale, shear and rotation transformations whereas the subsequent b-spline registration is a non-linear transformation. The initial affine step ensures that large architectural features in the image are registered before b-spline registers the finer cellular features. The affine and b-spline transformations are both tuned by a gradient-descent based optimiser to minimize the mutual information cost function.

Each affine and b-spline registration step incorporates a multi-resolution approach. The concept here is similar; to achieve better registration by registering large features before small features. At the beginning of the affine and b-spline step, a low-resolution image is used to encourage registration of the large features in the image. Gradually higher and higher resolutions are used to register every so finer features until the desired final resolution is reached. As registration is a computationally intensive process, especially for gigapixel WSIs, we registered smaller versions of the IHC and H&E images that were downscaled by 5 times - the downscale factor is user-adjustable. The final output of registration was colour 5x downscaled IHC images accurately registered to corresponding H&E images of identical size. As the H&E images may have captured a different field of view compared to the IHC images, any out of image pixels in the IHC images were filled in with white.

### Labelling images based on p53 staining

Registration transformed the p53 image to the same coordinate system as the corresponding H&E image. Thus, every pixel in the aligned p53 image referred to a pixel in the same location on the corresponding H&E image. This alignment was crucial for the p53 stain to accurately label the H&E image.

To label each pixel as one overlapping with cancer versus normal tissue, we applied thresholding to the p53 image. This process determined which pixels were positively (cancer) or negatively (normal) stained. The p53 IHC stain was visualized by the deposition of DAB on the tissue, giving positively stained tissue a brown colour. We distinguished DAB positive pixels, and hence p53 positive pixels, from the rest of the image by deconvoluting the RGB image into separate hematoxylin, eosin and DAB channels. This process was based on a method developed by Ruifrok and Johnston ^24^. In this way, we could focus our thresholding on the DAB stain, which reflects the level of p53 protein at each pixel.

We observed that the pixels within the DAB channel fell into three classes: p53 positive pixels; faint tissue background staining which we interpret as p53 negative staining; pixels of slide background where there is no tissue and no p53 stain. To simplify this into a two-class thresholding problem, we used the hematoxylin channel to separate the tissue from the slide background - we applied separate thresholding to the tissue only regions of the DAB channel. In both cases, we used Ostu thresholding which maximised the inter-class variance between two classes. Through segmenting the tissue with the hematoxylin channel, we distinguished the tissue by its low, but considerably greater than slide background, levels of stain. In addition, it ensured that we retained the nuclei which have high levels of hematoxylin and is where the p53 protein is localized. Following tissue thresholding, we applied the Otsu thresholding to only the tissue regions of the DAB channel and separated each pixel into two classes: a p53-positive class of high intensity pixels; a p53 negative class of low intensity background-stained pixels. This process was applied automatically and independently to each p53 slide so that pixel misclassification did not occur because of subtle differences in staining between p53 slides.

We split each H&E image into 224px x 224px tiles for model training and testing. Subsequently, we translated p53 pixel level classification to tile level cancer/normal classification. The registered p53 image was 5x down sampled to facilitate registration and it on this image that we determined pixel and tile labels, as it is aligned to the H&E. Thus, we analysed and labeled 5x down sampled tiles of 45px x 45px, of equivalent field-of-view to the original image. These tiles contain multiple cells – within a tumour infiltrated region of tissue, not all of these cells will be cancer. To ensure that we did not miss cancer cells while minimizing the levels of false staining, we labeled a tile cancer if more than 2% of the pixels within the tile were p53 positive. The remaining tissue tiles were labelled as normal or ‘non-cancer’.

In some cases, the p53 stain is not distinct enough to provide a definitive label to a tile so we label ambiguous tiles as uncertain and discard them. These ambiguous tiles may add noise to the training data and prevent accurate evaluation of the model’s performance. We addressed this issue by setting an upper and lower user-selectable DAB intensity thresholds to enable labelling of tiles as uncertain. These thresholds were applied to the mean DAB intensity of each tile. Tiles that that fell between these thresholds were labelled as uncertain and were not used for training or testing the model. The remaining cancer and non-tumour tile labels were transferred from the registered p53 image to the H&E tiles destined for model training.

### Splitting stained images into labelled tiles

We trained the model with 224px x 224px tiles from 10 H&E WSIs at 10x magnification. Due to tiling strategy, we could generate thousands of samples from each WSI which we pooled together for training the model. To safeguard against any registration errors and ensure accurate label transfer, if a p53/H&E pair of tiles had only one tile containing tissue, that H&E tile would be discarded. To assess a tile, we segment the tissue from the background in both p53 and H&E images using the GrabCut algorithm by Rother, et al. ^25^. In addition, to ensure a clean training dataset, only cancer-positive tiles from cancer samples were used and only cancer-negative tiles from the non-cancer samples were used.

### Training a convolutional neural network (CNN)

We used transfer learning to develop a VGG16 based CNN for classifying tiles as cancer or non-cancer. Our model utilized a VGG16 architecture and was pretrained on approximately 1.3 million images from ImageNet^26^, for feature extraction. By using weights pretrained on a large number of images, we can train our model a relatively small dataset and still achieve accurate predictions without overfitting. Features from each 224px x 224px tile were fed into a fully connected neural network to predict tile cancer status.

The complete CNN was trained on labeled H&E tiles generated from the 10 training WSIs at 10x magnification, for 100 epochs. We employed data augmentation to overcome overfitting and improve model generalizability. Since a given tissues extent of tumor cell infiltration remains the same regardless of the viewing angle or orientation, we randomly rotated and flipped tiles. The hyperparameters that performed best on the validation set were used for training the model that was used on all testing of unseen slides in this work. We implemented this system with Python using Tensorflow as the deep learning framework.

### Performance evaluations

We tested our model on H&E test slides, evaluating its performance compared to p53 stain patterns and pathologist annotations. We measured model performance by computing accuracy, confusion matrices and receiver-operating curves **(ROC)**. To evaluate performance against p53 annotations, we generated a test dataset using the same method described for the training dataset. Given that the sections had cellular mixtures, we generated tiles that solely represented cancer and normal tissues. For 13 of the 17 slides, we acquired pathologist cancer annotation drawings on the WSIs. We extracted the annotations and labeled tiles enclosed by the cancer annotation as cancer and labeled the remaining tissue tiles as non-cancer **(Supplementary Figure S4)**.

The main performance metrics are accuracy and ROC AUC. These are calculated by comparing the p53 and pathologist test dataset tiles labels with the labels predicted by our model **(Figure 4, 5 and Figure S5)**. Since cancer and non-cancer tiles do not evenly distribute in these datasets, we balanced the number of tiles for each class by subsampling the dominant class.

### TCGA validation

We validated our model on 24 colorectal cancer with H&E images. The data was obtained from the TCGA. We used our model predictions to estimate tumour purity and compared this to estimates of tumour purity derived from genome sequencing studies. For this image-based analysis, we calculated the proportion of the cancer tissue area to total tissue area by weighting tile predictions by the area of tissue within each tile. This is more accurate than using the proportion of cancer tiles to all tiles as some tiles, especially on the edge of the tissue. For example, a tile that is half background and half tissue would only contribute half a tile worth of area. We compare our estimate to seven method for determining tumor purity. This comparison included the programs ABSOLUTE^27^, EXPANDS^28^, ESTIMATE^29^, CPE^30^, InfiniumPurify^31^ and LUMP (leukocytes unmethylation for purity) **(Figure S6)**.

## RESULTS

### Molecular information for H&E images annotation

We developed a novel approach which leverages molecular annotations and deep learning methods to improve the identification of cancer cells **(Figure 1)**. The HEMnet development pipeline comprises four major steps: (1) data generation of paired P53 and H&E images, (2) preprocessing images and transferring of molecular label, (3) training neutral network, and (4) evaluating the performance of HEMnet **(Figure 1)**. The HEMnet pipeline was designed for applicability to any staining type or cancer type.

**Fig. 1.**
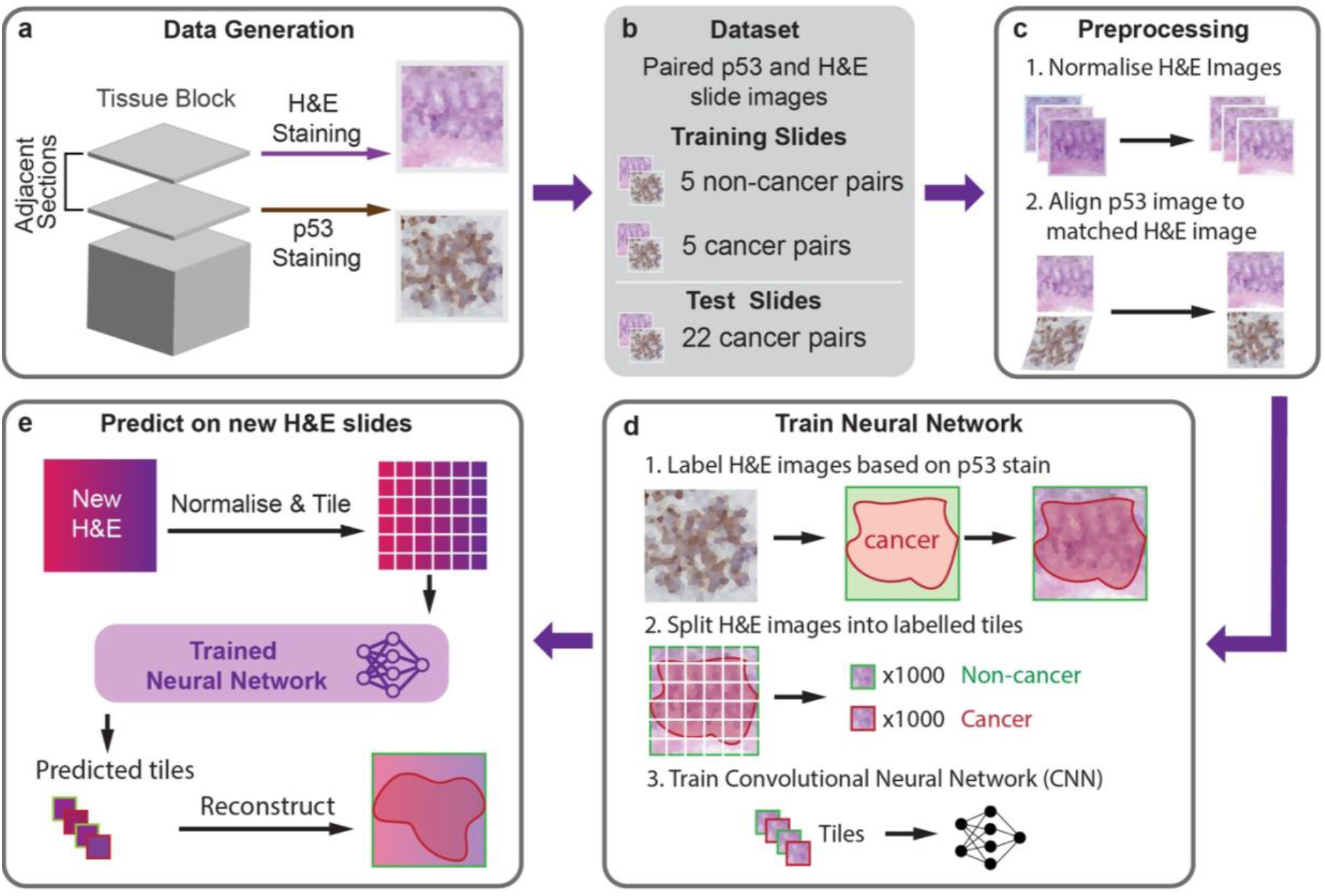
H&E Molecular neural network workflow overview. **a,** Matched p53 IHC stained and H&E stained WSI derived from two adjacent tissue sections. **b,** Training was performed on paired normal and cancer slides (five pairs). Test slides were held-back and are unseen. **c,** Preprocessing to account for technical variations in slide preparation through stain normalization and image registration **d,** Molecular labels were transferred from p53 to H&E images. Post label transferring, each image was tiled to generate thousands of small samples (224×224 pixels) to train a CNN **e,** Application of HEMnet to predict cancer from new clinical H&E images.

For this study, we developed HEMnet to identify tumor cells in H&E images of colorectal cancers. For step 1, we obtained 32 high-resolution H&E images and corresponding p53 IHC images from 27 cancer samples and 5 non-cancer samples. This was achieved by staining adjacent tissue sections with H&E and p53 to generate a matched paired WSIs for each tissue block. Step 2 is the novel contribution of HEMnet to transfer molecular labels to the H&E image. HEMnet takes advantage of molecular information, instead of manual pathologist annotations. We accomplished this by alignment of p53 molecular stained images to the corresponding H&E images at the pixel level **(Figure 3)**. The p53 stain pattern was, thereby, used to label cancer regions on the paired H&E images in an automated fashion, without the need for pathologist intervention. For step 3, each labelled H&E image was split into thousands of small tiles 224px x 224px so that from a small sample of 10 WSIs we can generate tens of thousands of training samples **(Figure 3d)**. We used these image tiles to train a deep transfer learning classifier to identify cancer regions in clinical H&E images using only tissue morphology features. Step 4 provides stringent validation criteria with independent datasets, comparing HEMnet with pathological annotation and with seven computational genomics diagnosis methods.

### H&E stain normalisation reduces colour variation

Besides realizing the novel concept of using molecular labels in deep learning model, the technical contribution of the HEMnet pipeline lies in the seamless pipeline, comprising a step to combine multiple images into a model training and testing dataset by normalizing different images, followed by fast and accurate label mapping, before training a neural network. Initially, WSIs with similar tissue structures stain different colours due to differences in slide processing (e.g. staining time, microscopy exposure). We address this issue with stain normalization, which caused these WSIs to take on the stain color profile of the template slide and increased the luminance to produce a white background **(Figure 2a-c, Figure S2)**. This method changed the mean R, G and B channel intensities of the normalized slide to closely resemble the template slide whilst retaining the R, G and B color distributions within the image. Across the 32 H&E WSIs, stain normalization reduced the variation in mean R, G and B channel intensities **(Figure 2d)**. In addition, it adjusted the median of the median channel intensities to move closer to the mean channel intensities of the template image. By normalizing all images before input into the model, we ensure the model can generalize to new slides stained differently to the training slides.

**Fig 2.**
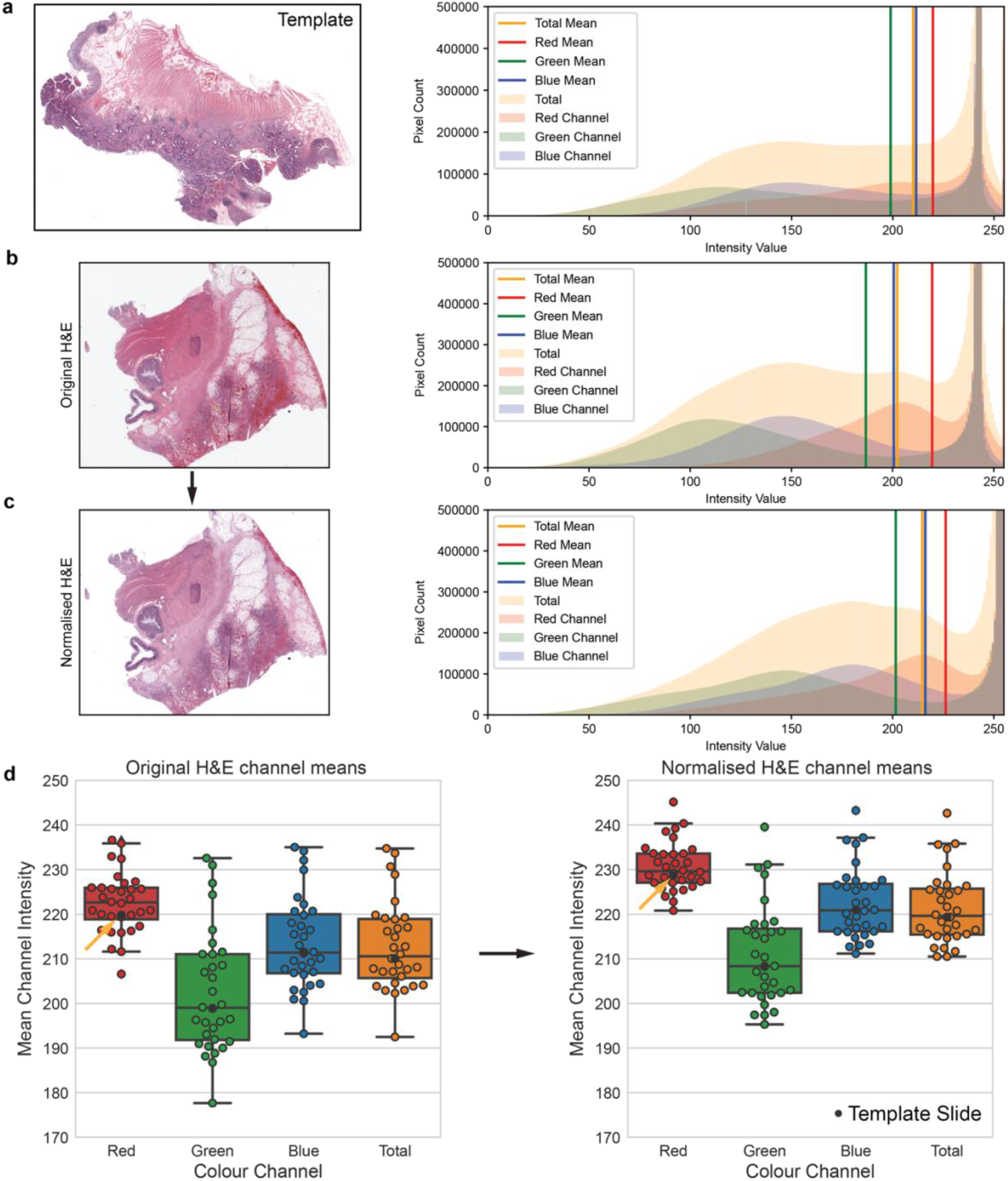
H&E stain normalization. **a,** Template slide – cancer slide with mean R, G and B channel intensities most similar to the median of the mean channel intensities of all images. Histograms shown for 2x magnification image (**a, b, c**) **b,** H&E image before normalization **c,** H&E image after normalization more closely resembles template image. Image brightness is increased and pixel intensity distributions are retained. **d,** Normalization of all slides (n=32). Reduced variation of mean channel intensities after normalization was observed. Template slide mean channel intensities are closer to the median after normalization (indicated by arrows) and interquartile range was shrunken.

### Transferring p53 molecular labeling to corresponding H&E images

The WSIs from corresponding p53 and H&E-stained slides often were misaligned **(Figure 3a)**. For the p53 positive cells to accurately map to cancer cells on the H&E images, we realigned p53 images to their corresponding H&E images though HEMnet automated image registration **(Figure 3c)**. Our intensity-based registration approach was fast and accurate as we optimized mutual information **(Figure 3b, c)**. Next, we labelled the H&E image based on the p53 staining pattern where p53 positive regions are labelled as cancer, vice versa. To counteract limitations of p53 staining in marking cancer cells, only p53 positive tiles from cancer slides and only p53 negative tiles from non-cancer slides were used for training. All the other tiles were labelled as uncertain and excluded from any additional processing. At x10 magnification, a single WSI can generate thousands of tiles for training **(Figure 3c)**. We generated 224×224 pixel tiles from the molecular labelled H&E images to train a VGG16 deep learning model **(Figure 3d)**.

**Fig 3.**
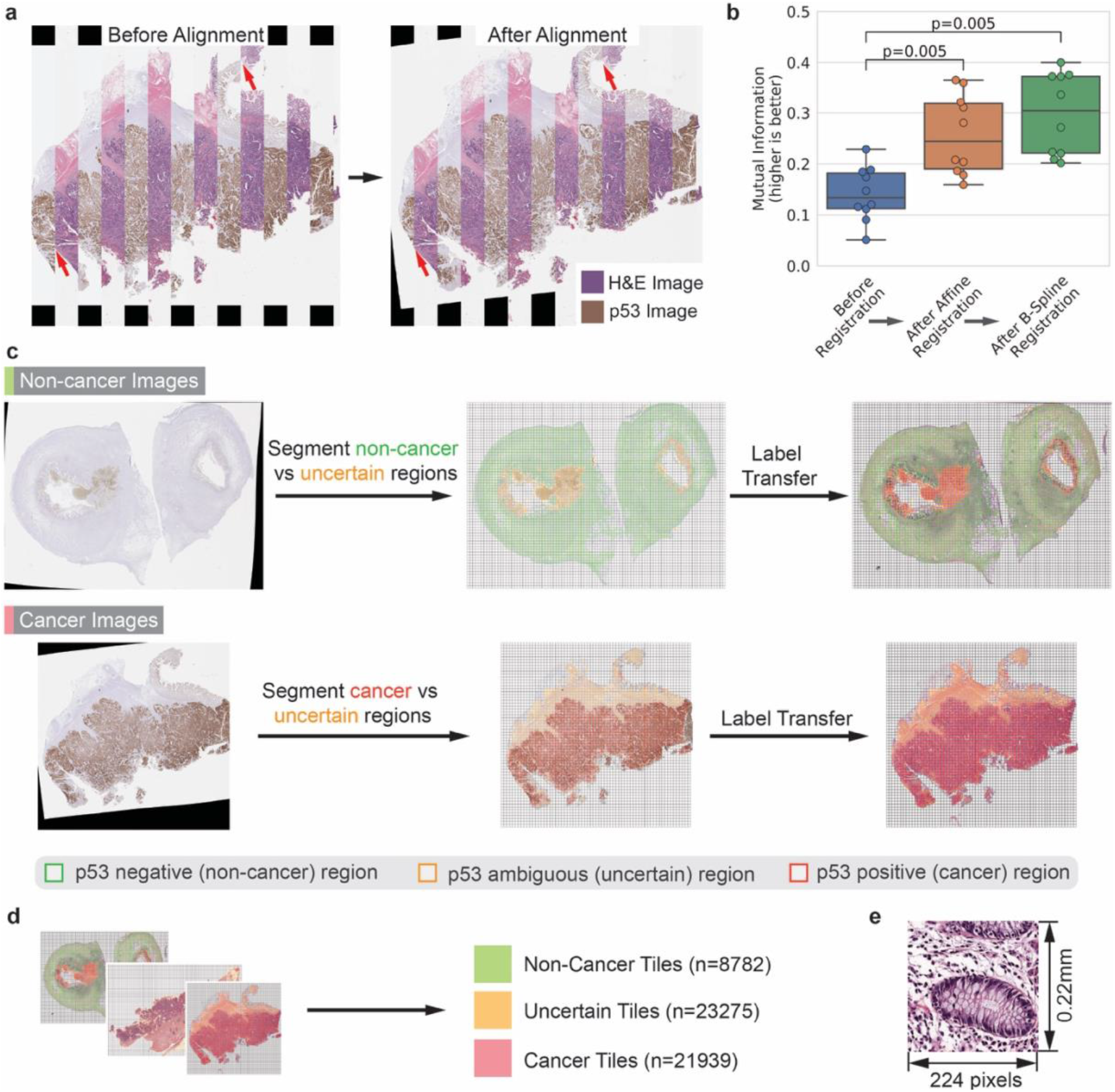
Molecular labelling of H&E images to train Neural Network. **a,** Overlay of H&E and matched p53 image showing improved alignment after registration, highlighted by red arrows **b,** Accurate alignment of p53 images to corresponding H&E images. Successive affine and b-spline registration increases mutual information, a measure of image similarity. Significance testing with T-test. **c,** Segmentation of p53 images to label matched H&E images where only non-cancer tiles are generated from non-cancer slides, vice versa **d,** 10 training H&E images generated tens of thousands of tiles, increasing sample size **e,** Example of cancer tile generated at x10 magnification and used for training neural network.

### Molecular annotation quality control produces a high-confidence dataset

The *TP53* tumour suppressor gene is the most commonly mutated gene in human cancers (50%) and disproportionately has mutations and other genetic alterations for up to 70%-80% of colon cancers^32,33^. As a result of its general prevalence, it provides a highly generalizable way to molecular annotate a broad range of cancers. Similar to other IHC markers, p53 staining has its limitations as within one image or between images, the marker is not always indicative of cancer, vice versa. For example, overexpression and positive staining for p53 may occur in normal cells responding to DNA damage. In addition p53 may be absent in cancer cells with *TP53* gene deletions^14^. To overcome these limitations, when training our model, we only considered p53 positive cells as cancer if they come from a cancer slide and only p53 negative cells from slides where the cells have a normal morphology **(Figure 3d)**. In this way, we were confident that cells were correctly labelled, with 8,782 non-cancer tiles and 21,939 cancer tiles. We removed 23,275 tiles that had some levels of uncertainty **(Figure 3d)**.

### High performance automated assessment of cancer cell abundance and spatial distribution

We applied the trained HEMnet to unseen WSIs to predict cancer regions. Of the 17 unseen H&E slides in the test dataset, all had corresponding p53 stained slides and 13 had additional pathologist annotation of the cancer region. We found that HEMnet could accurately predict p53 stain pattern (ROC AUC = 0.73) and pathologist annotated cancer regions (ROC AUC = 0.84), (**Figure 4a,b**). These results suggest that p53 positive cancer regions for a given tissue sample can be predicted from its general morphology using a classifier developed with molecular labelled H&E images.

**Fig 4.**
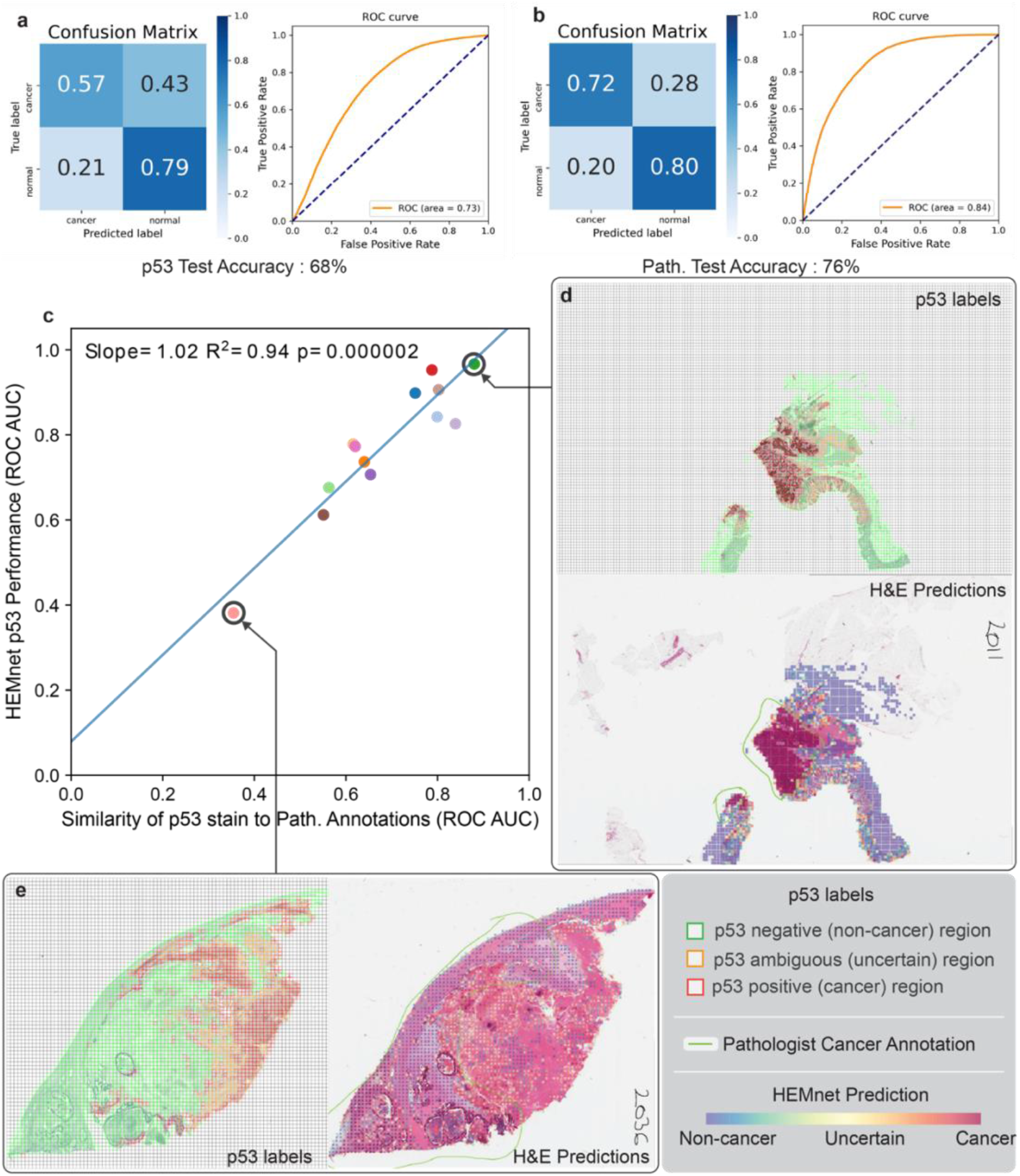
HEMnet performance on unseen H&E slides. **a,** Prediction of p53 stain pattern on 17 unseen H&E slides **b,** Prediction of cancer regions on 13 unseen H&E slides compared to pathologist annotations **c,** Prediction performance of p53 stain pattern is positively correlated with the ability of p53 to mark cancer regions on the tissue, as annotated by pathologist. **d,** HEMnet accurately predicts cancer regions annotated by pathologist (bottom) and p53 stain (top) when p53 stain agrees with pathologist annotation **e,** HEMnet predicts cancer regions even when p53 stain pattern (left) disagrees with pathologist ground truth annotations (right).

Comparing the p53 labeled tiles to pathologist labelled tiles from the same location, we found an overall agreement in tile labels (ROC AUC = 0.67) **(Supplementary Figure S6)**. However, this agreement was not absolutely perfect. To evaluate any discrepancies, for each slide we measured the ability of p53 stain to annotate cancer. This analysis involved calculating the ROC AUC between p53 stain and ground truth labels of tiles per a pathologist. We found that HEMnet p53 performance (ROC AUC) was higher in slides where p53 more accurately labelled cancer (p53 vs pathologist tile labels ROC AUC) with a significant correlation as noted by a Pearson coefficient of 1.02, and R^2^=0.94 **(Figure 4c)**. This result indicated that the model learnt to recognize specific morphology features of cancer cells and was not strictly limited to identifying cells with high levels of p53. This likely because cancer cells are morphologically distinct from normal cells whereas the differences in morphology between p53 positive and negative cells are more subtle. We noted that there were examples demonstrating that HEMnet can identify the cancer marked by the pathologist, even where the cancer is not identified by the p53 stain **(Figure 4d, e)**. Overall, the results suggest that HEMnet is able to accurately identify tissue morphology features of cancer.

### External validation and application to TCGA suggests the broad applicability

As an independent validation using an external dataset, we applied HEMnet to colon adenocarcinoma samples from TCGA colon cancer samples to investigate the generalizability and clinical application of the method (Table S1). We used an unmodified HEMnet model trained by the in-house dataset described in this study to predict on H&E WSIs of colon adenocarcinoma. By combining the tile level prediction with the cellular content of each tile, we calculated the proportion of cancer tissue to total tissue for each slide (Table S1, Figure 5a). This acts an approximation of tumour purity which we compared to sequencing method estimates from matched genomic data. There are several differences between our colon cancer data and the TCGA data. Most importantly, the sequencing was not performed on the same tissue used for diagnostic imaging. Despite these challenges, we found a significant correlation between our method and tumour purity as estimated by ABSOLUTE, with a regression coefficient of 0.8, as shown in Figure 5. Furthermore, we found that HEMnet performs well regardless of the TP53 mutation background (Figure 5a). This analysis suggests that HEMnet can generalize to new colorectal clinical data and is able to reliably predict on TCGA images.

**Fig 5.**
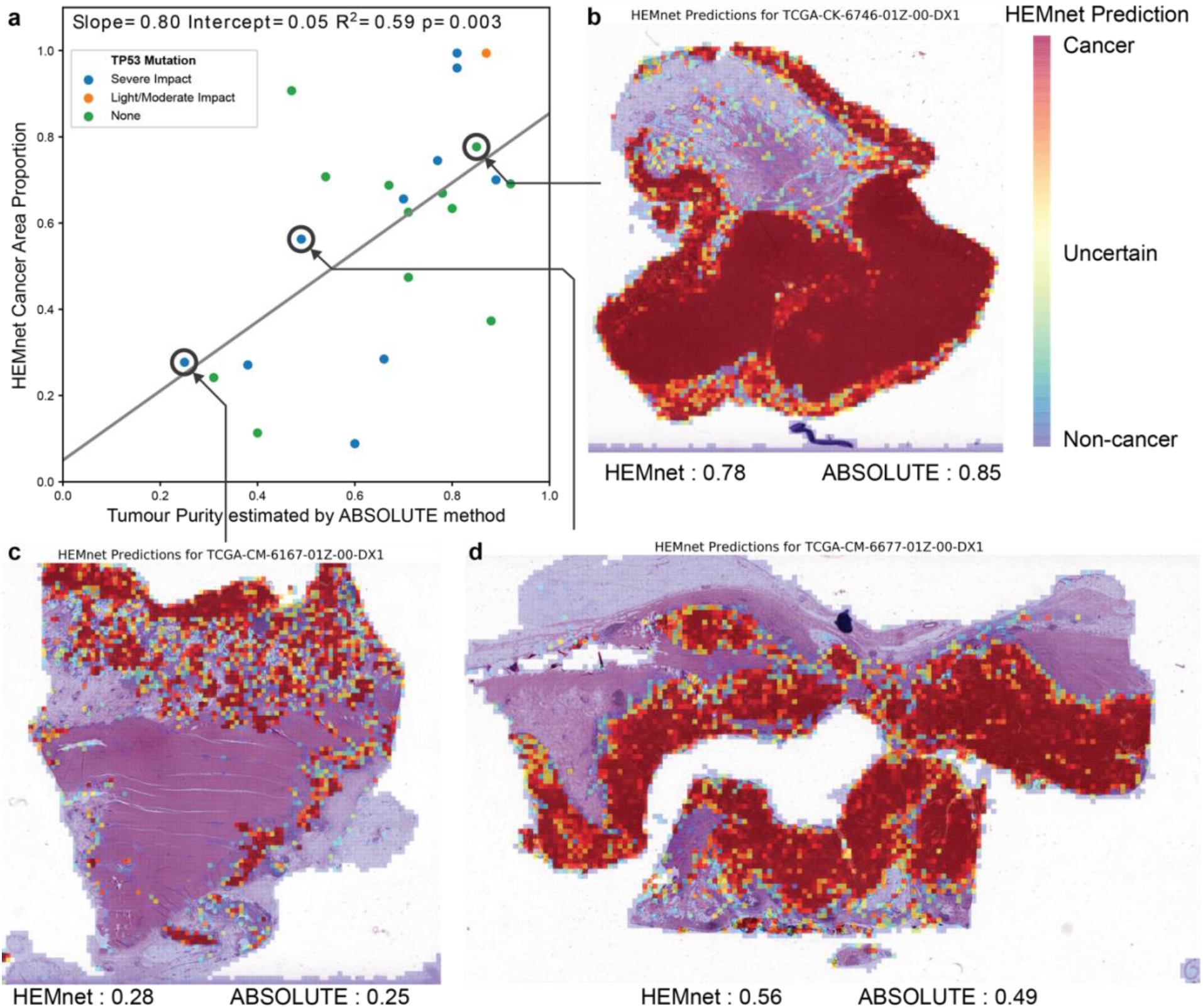
External Validation on The Cancer Genome Atlas (TCGA) **a,** Comparison of HEMnet estimation of tumour purity – approximated by the proportion of cancer tissue area to total tissue area – to sequencing estimates of tumour purity using the ABSOLUTE method (n=24). The colours of the dots represent three categories of TP53 mutations from the TCGA data. **b, c, d,** HEMnet cancer predictions on formalin fixed TCGA slides for low (c), medium (b) and high (d) tumour purity colon adenocarcinoma.

## DISCUSSION

Histopathological examination of H&E images has been the gold standard for pathologic diagnosis of almost all suspected cancer patients^3,34^. Modern applications of machine learning tools to analyse H&E images have been flourishing recently^6,35^, with some of the computer-assisted image diagnosis tools already approved by the Food and Drug Administration (FDA)^36^. Hundreds of deep learning produced have made methods available for using just H&E images to detect and diagnose cancer^6^. Although some of these methods have achieved high performance, they all rely on pathological annotation for labelling/segmenting images into multiple tissue regional classes^6,37^. They also require a large number of annotated images for model training and evaluation^38,39^ and the lack of large annotated datasets is a major challenge for deep learning image analysis^6^. We developed HEMnet as a novel cancer diagnosis framework that uses digital labelling and neural network to address these challenges.

HEMnet combines two common types of histopathological WSI data, namely H&E staining and immunohistochemistry staining images. The novelty in HEMnet pipeline lies in the molecular label transferring, which allows for the use of pixel-level molecular information cancer cells (e.g. P53 positive/negative pixel), with thousands time higher resolution than manual pathological segmentation. In HEMnet, we solved several key technical challenges to allow for accurate, fast and generalizable label transferring, with the ultimate aim that HEMnet can be implementable to different datasets, including those with a high level of technical variation. Briefly, technical variation is introduced by the tissue sectioning, mounting, staining and imaging processes. Very few studies investigated the intrinsic technical variations, like contrast, brightness, or signal to noise ^6^. Different to most methods, HEMnet implements an optimized pipeline for preprocessing, allowing removal of technical variation between images. HEMnet include functionalities to thoroughly perform background correction, normalization, alignment, registration, and label transferring. Prior to normalisation, luminosity standardization was performed to correct for image brightness. We compared three normalisation methods, Vahadane^19^, Reinhard^21^ and Macenko^22^, and confirmed the better performance of the Vahadane method (set as default). The image registration implements a probabilistic approach with mutual information maximization. We compared multiple options and found that intensity-based registration, and the sequential combination of Affine followed by B-spline registration^40^, using a gradient-descent based optimizer to minimize mutual information loss perform well for registering H&E image data. We also assessed the computation and running time, as registration is an intensive process. Down-scaling was found as a practical solution. Finally, to label the registered image, we developed a tile-level thresholding strategy to distinguish cancer, non-cancer and uncertain labels for every tile of 224px * 224 px. The tile-level labelling with thresholding, categorizing and filtering steps allows us to create a high-quality training (and evaluation) data set for neural network, minimizing the technical noise from registration errors and uncertain labelling.

Overall, the label-transferring solution implemented in HEMnet represent a significant technical advance and is needed to the increasingly important digital histopathological analysis field. The label transferring brings about three key beneficial effects on model training. First, the pixel-level labels allow us to divide one image into hundreds to thousands of smaller, high-resolution, molecular labelled tiles, thereby increasing sample sizes for model training and testing. This enables development of accurate models with few slides, unlike existing methods which require a thousands of whole slide images ^6,41^. In general, tiling of WSI yields the large amount of data for training neural network, thus would be robust to incomplete molecular markers. It was demonstrated by the fact that HEMnet successfully identified some non p53 stained cells as cancer cells (Figure 4). With pixel-level labelling, the classification of cancer cells is at hundreds to thousands of times higher resolution than macroscopic drawings by pathologists. Moreover, molecular labelling is automated, making the output less dependent on the laborious, manual and variable annotations by trained pathologists.

HEMnet, with its novel label transferring approaches, can be beneficial for a large range of applications. When processing an independent validation set not used in the original learning process, HEMnet predicted the same overlapping region delineated though a pathology annotation (ROC AUC = 0.84). We validated HEMnet by systematically comparing HEMnet with other methods and with the ground truth pathological annotation and found highly correlated results with other independent methods (correlation coefficient in predicting cancer purity = 0.8) using TCGA dataset^42^. The generalization to other types of markers and cancer, for example HER2 for breast cancer, is possible with further validation. The feasibility of correlating H&E images with IHC image by deep neural networks has been investigated for the case of SOX10 staining^43^ and fluorescent cancer marker images like pan-cytokeratin (panCK), or α-smooth muscle actin (α-SMA)^44^. HEMnet was developed using p53 IHC staining as an appropriate colorectal cancer marker that is expressed in 70%-80% of colon cancers^12^. We expect that the HEMnet label transferring and thresholding approaches to define positive cancer labels can be generalized to other cancer types and immunohistochemistry markers. We expect that HEMnet can be readily adaptable to training new data as the design of the framework take into account technical variation and scalability as discussed above and as confirmed by the test on the TCGA dataset robust performance. The novel label transferring pipeline can be expanded to many other applications to integrate imaging data from adjacent tissue sections. We made HEMnet an easily adaptable tool for most users through the interactive Google Colaboratory workspace, which allows users to upload their data and use our pretrained model for neural network prediction.

## CONCLUSIONS

HEMnet is currently the unique molecular modelling approach that utilizes both H&E and IHC images for quantitatively classifying cancer cells within tissue sections. We expect that HEMnet has the potential to be used as a computer-assisted tool that help pathologists by suggesting important regions, such as cancer parts, in the tissue ^35,45^. HEMnet does not require human pathological annotation, automatically labelling images at pixel resolution. The application of software like HEMnet can benefit cancer diagnosis by unprecedented resolution, efficiency, reproducibility, accuracy, speed, reduced cost and increased access to pathological services. In an aging society where more biopsies are available while there is a lack of professional anatomic pathologists ^46^, such computational innovation is increasingly important. We believe HEMnet can further accelerate computational pathology application and integration into the pathology workflow routine, assisting in disease diagnosis and ultimately removing missed diagnosis and improving patient outcomes. We provide HEMnet as an open-source software and also as an accessible cloud-based prediction tool that allow users to analyse their images without a requirement for further programming.

## Availability of Data and Materials

The datasets used and/or analysed during the current study are available from the https://dna-discovery.stanford.edu/research/web-resources/HEMnet. The source code, tutorials and interactive analysis tools are available at https://github.com/BiomedicalMachineLearning/HEMnet. We also provide cloud-based implementation of the HEMnet **(Figure S7),** available as Google Colab notebook and an ImJoy application (links to these apps are on HEMnet github page). HEMnet is also available as an open-source PyPI python package (https://pypi.org/project/hemnet).

## Authors’ contributions

Q.N., A.S., N.A., HJ.L., X.T. and H.P.J. conceived experiments and developed the algorithms. A.S., X.T., wrote the software. A.S., HJ.L., X.T. and Q.N. conducted experiments and analysed data. A.S., HJ.L., Q.N., and H.P.J., wrote the manuscript. All authors have reviewed and approved the manuscript.

## ACKNOWLEDGEMENT

We thank all members in Nguyen’s Genomics and Machine Learning Lab and the Ji Research Group helpful discussion. We appreciated initial work on the alignment between H&E and IHC by Aykan Ozturk.

