## SupplementaryFigures and Table for "A Deep Learning Model for Molecular Label Transfer that Enables Cancer Cell Identification from Histopathology Images"

### SUPPLEMENTARY INFORMATION

#### Supplemental Figures:

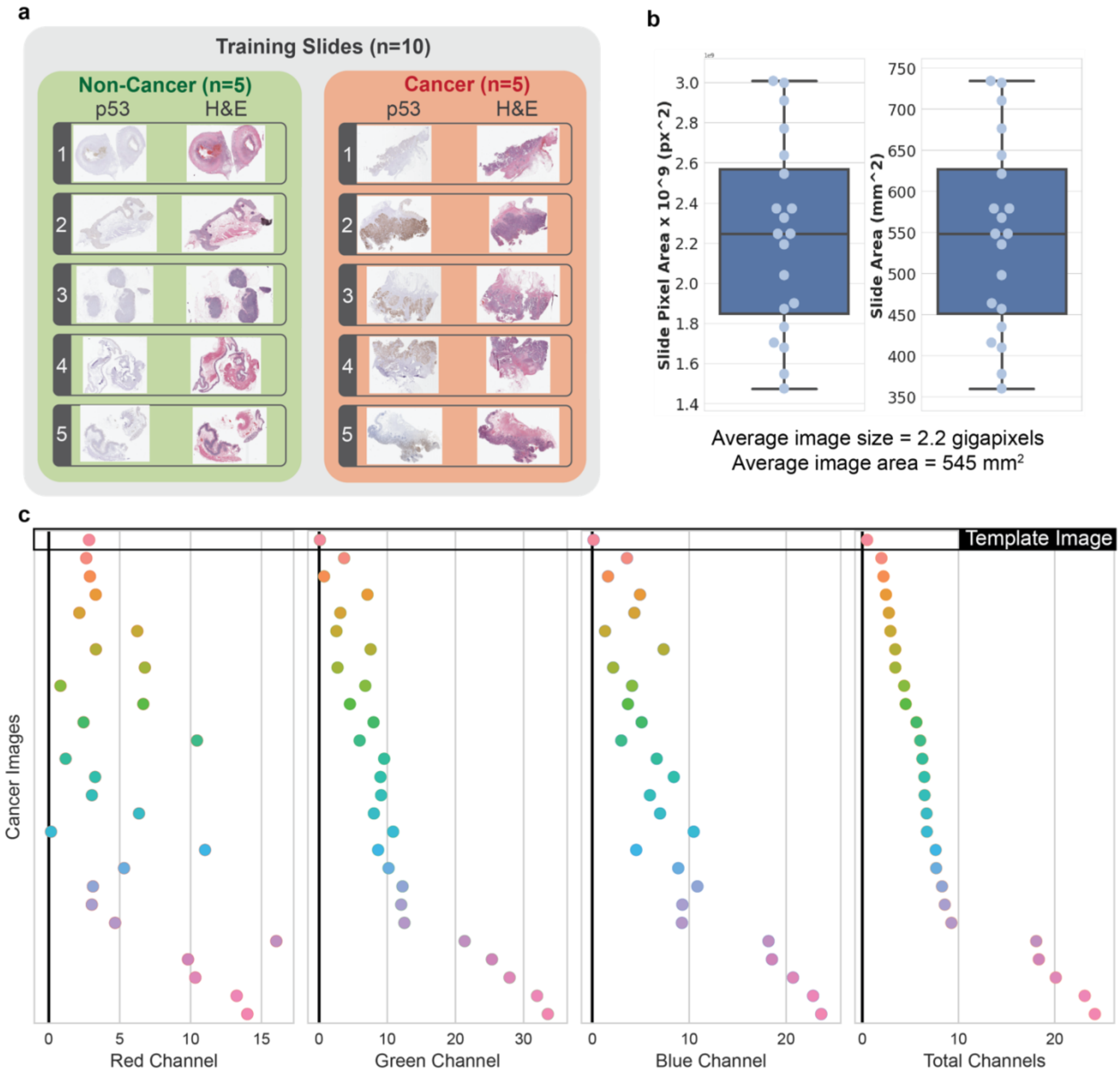


Supplemental Fig. S1. **Overview of Dataset a,** Training slides containing paired H&E and p53 slides – 5 pairs cancer, 5 pairs non-cancer. **b,** Distribution of training slide pixel areas in gigapixels and slide areas in mm^2. **c,** Comparison of the mean R, G, B and total channel intensities of cancer images with the median channel intensities of all images. Each subplot shows the absolute difference between the mean intensities of cancer images to the median intensity of all images, for a particular channel. The cancer image most similar to the median of all images is selected as the template image against which all other images will be normalized against.

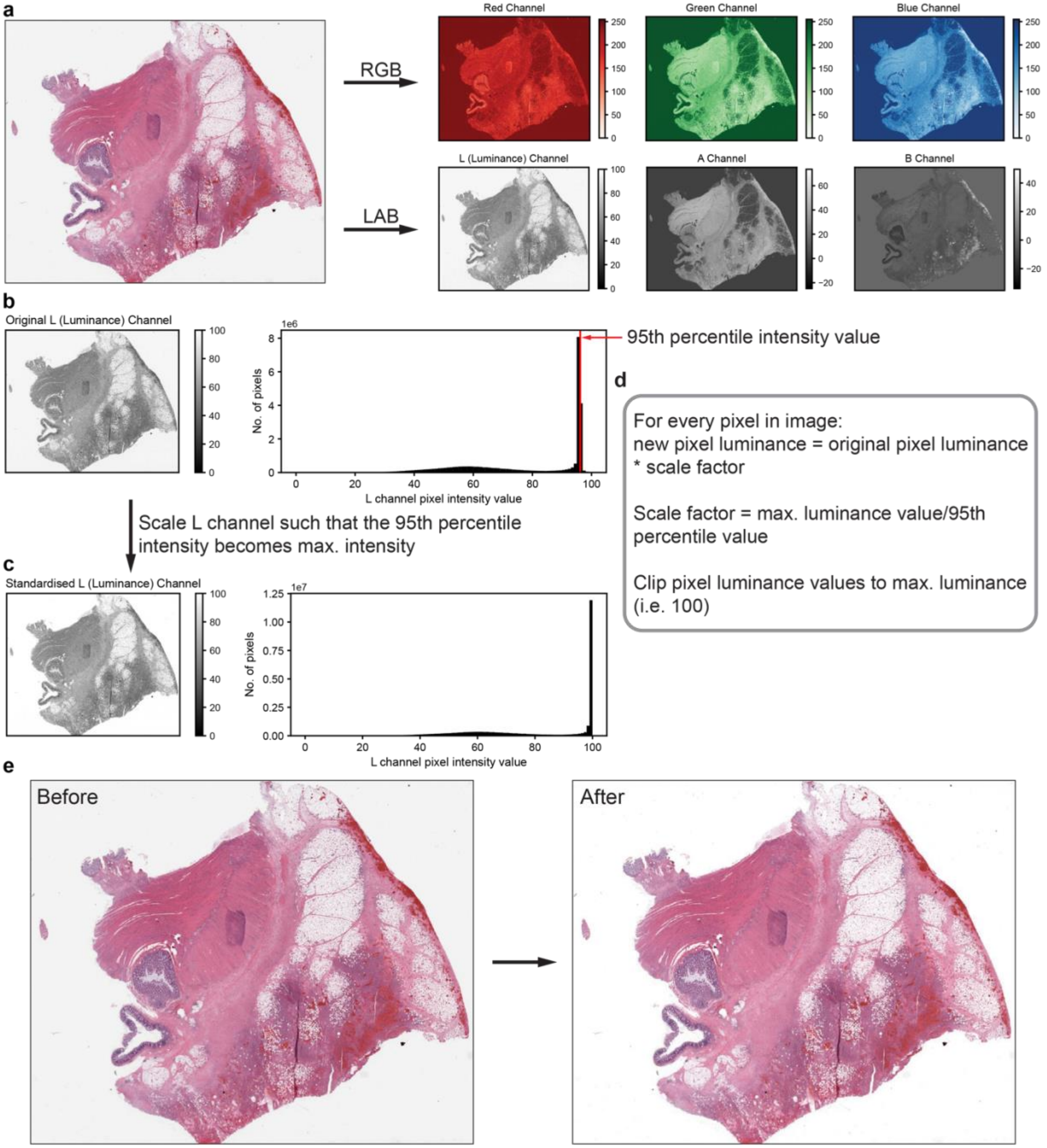


Supplemental Fig. S2. **Luminance Standardization workflow a,** Conversion of H&E WSI from RGB colourspace to LAB colourspace. **b,** Luminance (L) channel of WSI showing that few pixels are at 100 intensity, indicating the background is gray and not white. Histogram shown for 2x mag. image (**b, c**) **c,** Scaling of luminance (L) channel intensity so that 5% of the slide is at max luminance – this makes the background white. **d,** Pseudocode concept of luminance standardization as applied to each pixel **e,** Before and after standardization showing transformation of background from gray to white.

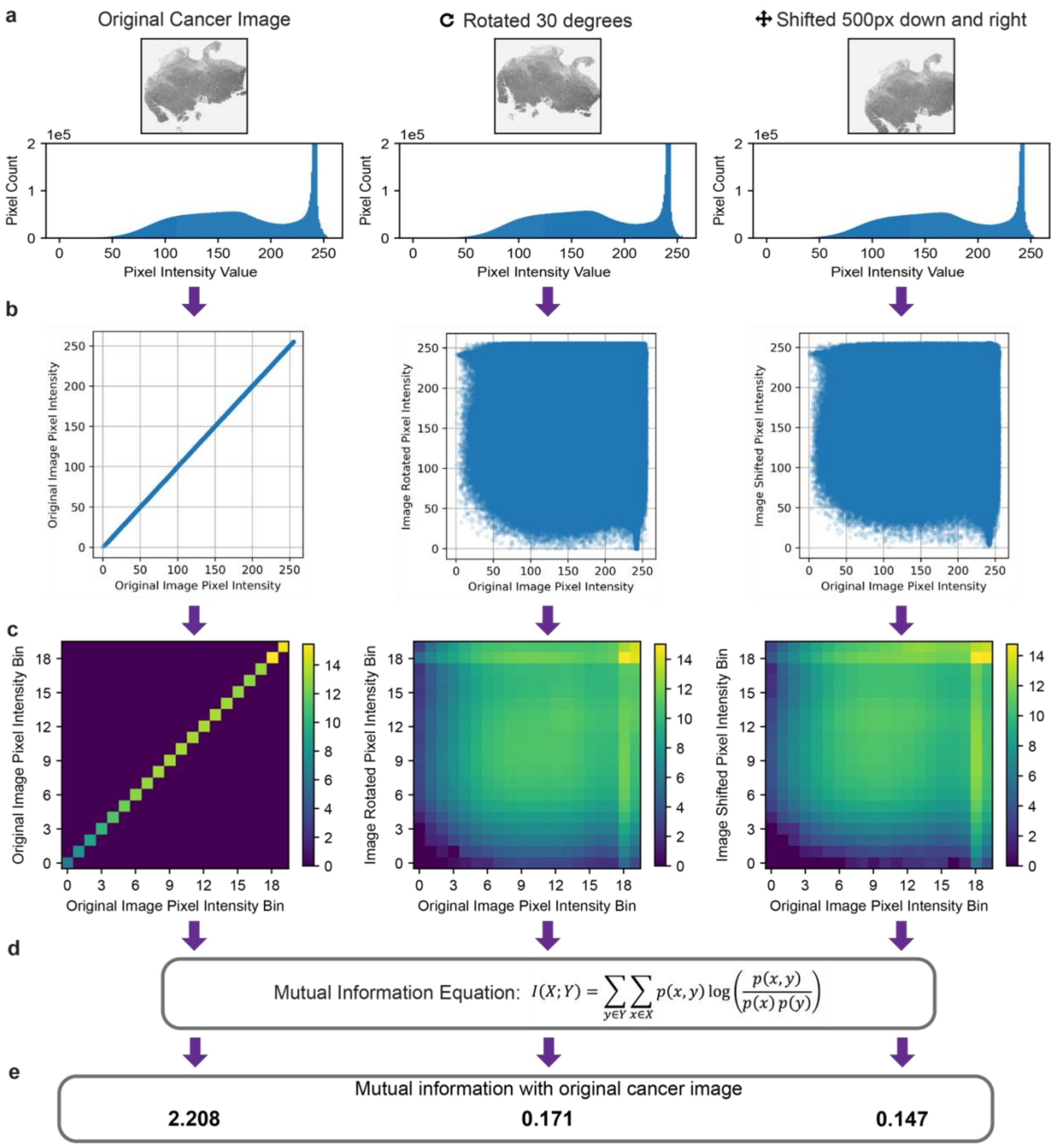


Supplemental Fig. S3. **Mutual information measures image similarity**. Demonstration of mutual information calculation on images**. a,** Grayscale WSI images at 2x magnification with different transformations yet similar pixel intensity distributions. **b,** Visualization of joint probability distributions. **c,** Log scaled 2D heatmaps of joint probability distribution. **d,** Mutual information calculated using marginal and joint probability images **e,** Mutual information between transformed images and original cancer image showing that mutual information is higher in aligned images.

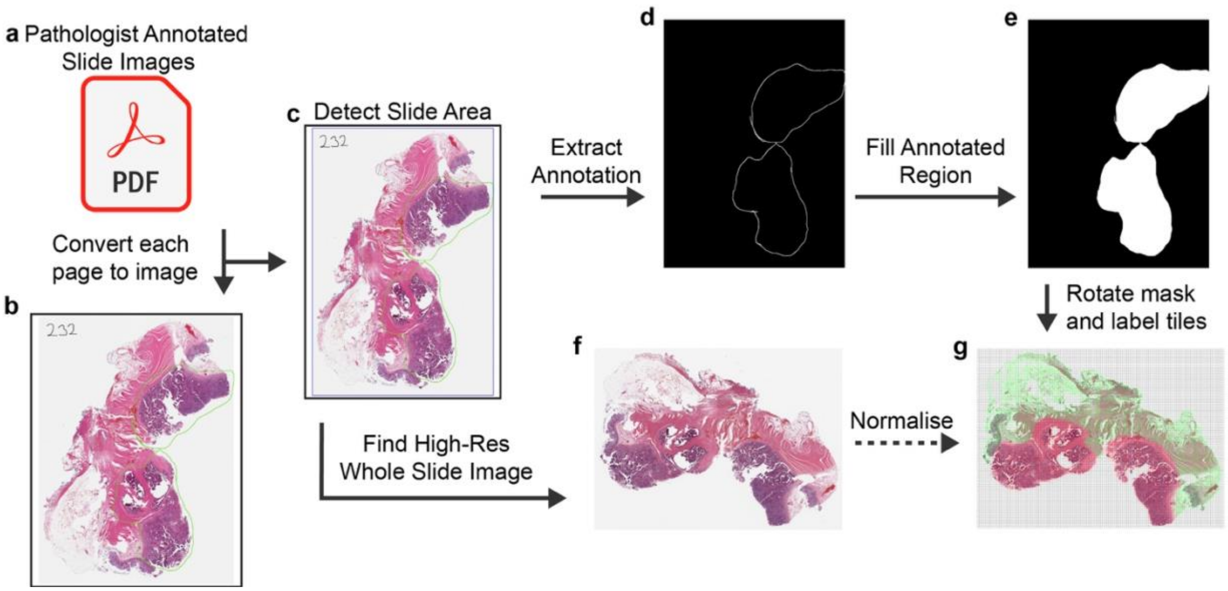


Supplemental Fig. S4. **Extraction of Pathologist Annotations a,** PDF file containing pathologist annotations on low-resolution slide images. **b,** A single page with green pathologist annotations on a slide image. **c,** Detection of the bounds of slide image to remove white borders. **d,** Extraction of green cancer annotation line as a mask. **e,** Flood fill of annotation regions to form a pixel level cancer mask, upscaled to match size of high-resolution WSI. **f,** Identification of high-resolution WSI from low-resolution pdf image. **g,** Normalised and tiled WSI with each tile labelled as cancer (red) or non-cancer (green) based on pathologist cancer annotation mask.

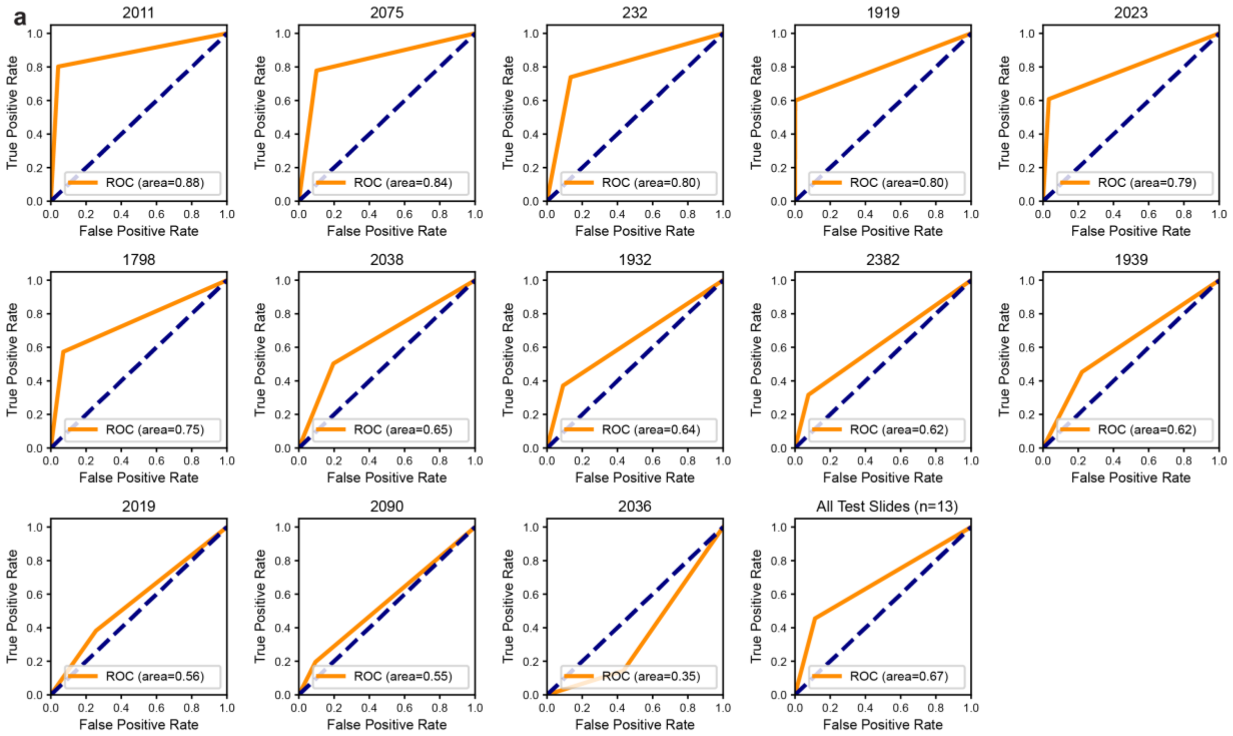


Supplemental Fig. S5. **p53 vs Pathologist annotations a,** Receiver operating curves (ROC) for test slides tiles with labeled by p53 and Pathologist annotations (n=13) showing that p53 annotations often, but not always, agree with pathologist annotations. Each plot shows the ROC for tiles from a particular test slide, expect the last plot which shows the ROC for all test slides. Higher ROC area under the curve indicates more tiles labeled by p53 staining had the same cancer/non-cancer labels as tiles of the same location labeled by pathologist annotation.

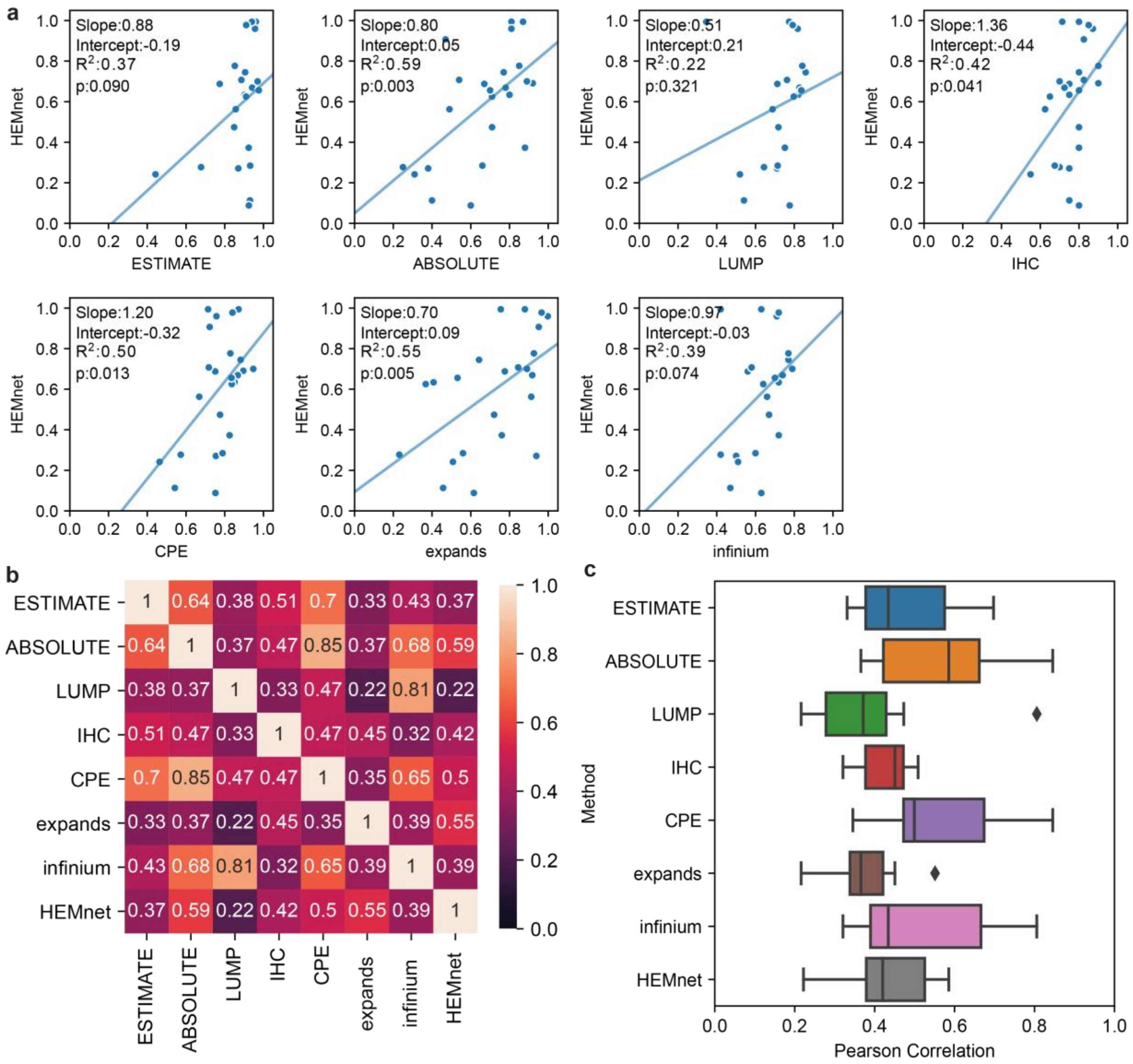


Supplemental Fig. S6. **HEMnet vs sequencing estimates of tumour purity a,** Individual plots of HEMnet cancer area proportion vs different sequencing estimates of tumour purity. **b,** Pearson correlations between different estimates of tumour purity. **c,** Boxplot of Pearson correlations between one method against all other methods. HEMnet performs similarly to other methods.

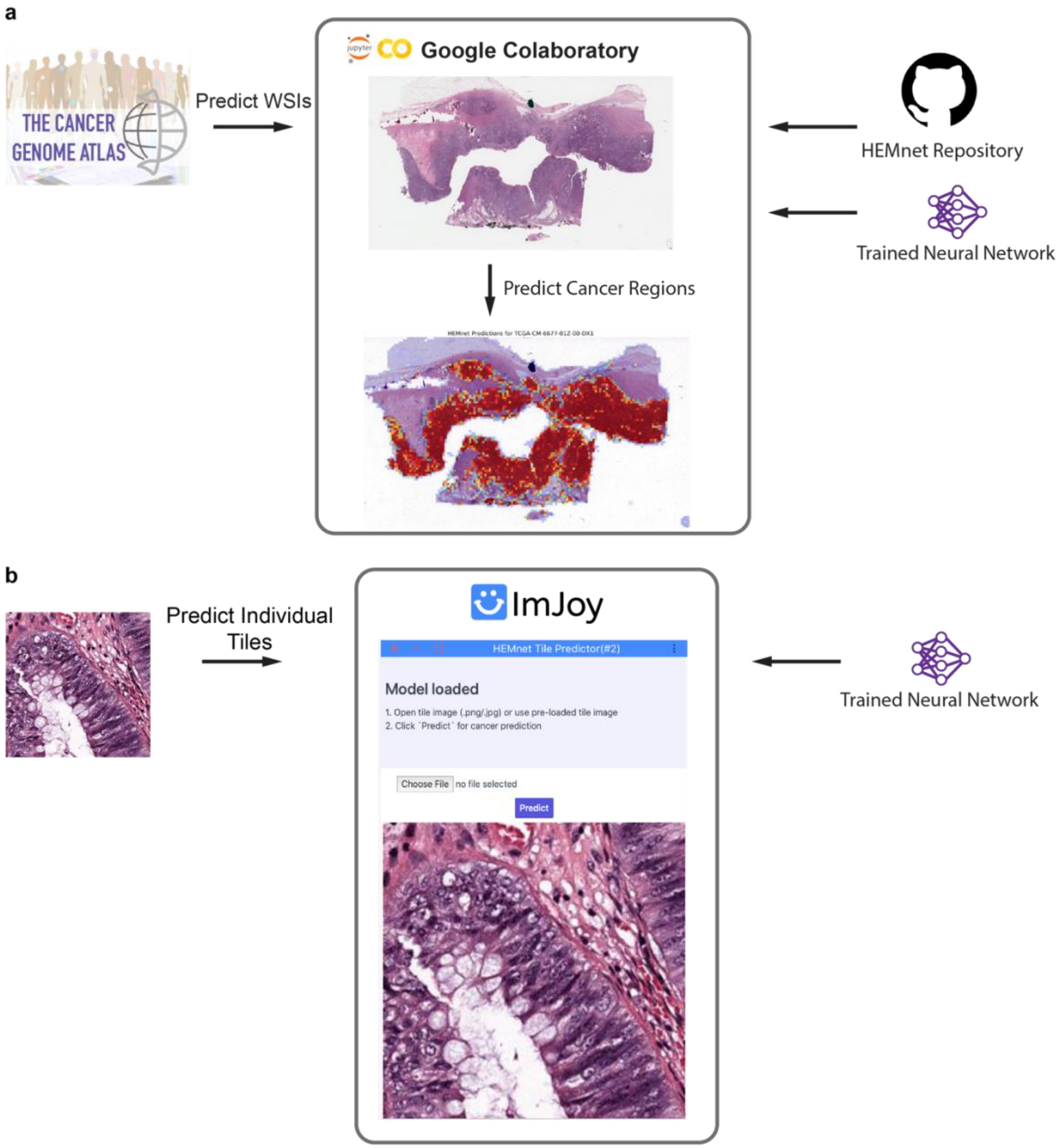


Supplemental Fig. S7. **HEMnet Demos in the cloud a,** Google colaboratory notebook for using HEMnet and our trained colorectal cancer model to predict on slides from TCGA. Outputs predicted cancer regions and estimated cancer area proportion **b,** ImJoy plugin for predicting on individual tiles using our trained colorectal cancer model.

**Supplemental Table:**

Table S1. Validation results using TCGA Colon Adenocarcinoma (COAD) dataset.

| Sample IDs | WSI Area (mm^2) | Tissue Area (mm^2) | Total Tiles | Cancer Tiles | HEMnet* Cancer Tile Proportion | HEMnet Cancer Area Proportion | ESTIMATE* | ABSOLUTE* | LUMP* | IHC* | CPE* | EXPANDS* | infinium* | TP53 Mutation Type |
|---|---|---|---|---|---|---|---|---|---|---|---|---|---|---|
| A6-5656 | 261.26 | 18.58 | 529 | 470 | 0.888 | 0.994 | 0.9606 | 0.81 | 0.774 | 0.714 | 0.714 | 0.879 | 0.63 | Missense_Mutation |
| A6-6650 | 448.67 | 76.96 | 1901 | 1633 | 0.859 | 0.960 | 0.9562 | 0.81 | 0.818 | 0.871 | 0.757 | 0.998 | 0.71 | Missense_Mutation |
| AA-3715 | 318.48 | 74.11 | 7886 | 6533 | 0.828 | 0.907 | NA | 0.47 | NA | 0.825 | 0.722 | 0.951 | NA | None |
| AA-3973 | 127.58 | 44.20 | 4919 | 3363 | 0.684 | 0.691 | NA | 0.92 | NA | 0.900 | 0.896 | 0.898 | NA | None |
| AD-6889 | 220.43 | 121.54 | 3032 | 2764 | 0.912 | 0.994 | 0.9399 | 0.87 | 0.346 | 0.800 | 0.871 | 0.755 | 0.42 | 3'UTR |
| AZ-4615 | 560.41 | 235.96 | 6421 | 3640 | 0.567 | 0.688 | 0.7739 | 0.67 | 0.713 | 0.750 | 0.751 | 0.776 | 0.56 | None |
| AZ-4682 | 539.76 | 174.56 | 5569 | 2602 | 0.467 | 0.669 | 0.9415 | 0.78 | 0.827 | 0.725 | 0.867 | 0.918 | 0.74 | None |
| AZ-5403 | 852.98 | 638.19 | 13865 | 3567 | 0.257 | 0.271 | 0.87 | 0.38 | 0.711 | 0.750 | 0.754 | 0.939 | 0.5 | Missense_Mutation |
| CA-6715 | 395.61 | 232.72 | 5840 | 3560 | 0.610 | 0.700 | 0.9696 | 0.89 | 0.762 | 0.700 | 0.947 | 0.890 | 0.79 | Missense_Mutation |
| CK-5913 | 488.02 | 291.19 | 7160 | 3044 | 0.425 | 0.474 | 0.8486 | 0.71 | 0.718 | 0.800 | 0.776 | 0.721 | 0.67 | None |
| CK-5914 | 504.17 | 221.81 | 5200 | 3705 | 0.713 | 0.745 | 0.9052 | 0.77 | 0.859 | 0.800 | 0.882 | 0.643 | 0.77 | Missense_Mutation |
| CK-6746 | 436.20 | 232.33 | 5409 | 3753 | 0.694 | 0.777 | 0.852 | 0.85 | 0.842 | 0.900 | 0.829 | 0.926 | 0.77 | None |
| CK-6747 | 580.80 | 265.62 | 6853 | 3680 | 0.537 | 0.634 | 0.9021 | 0.8 | 0.821 | 0.750 | 0.851 | 0.408 | 0.72 | None |
| CM-5344 | 546.99 | 312.38 | 7301 | 4771 | 0.653 | 0.707 | 0.8858 | 0.54 | 0.763 | 0.825 | 0.718 | 0.845 | 0.58 | None |
| CM-6167 | 472.38 | 261.19 | 6217 | 1567 | 0.252 | 0.277 | 0.6777 | 0.25 | 0.644 | 0.700 | 0.573 | 0.231 | 0.42 | Missense_Mutation |
| CM-6171 | 863.72 | 294.10 | 7046 | 6462 | 0.917 | 0.978 | 0.9107 | NA | 0.792 | 0.850 | 0.840 | 0.966 | 0.72 | None |
| CM-6677 | 678.28 | 292.76 | 7136 | 3443 | 0.482 | 0.563 | 0.8565 | 0.49 | 0.688 | 0.625 | 0.668 | 0.912 | 0.66 | Missense_Mutation |
| D5-5537 | 953.17 | 240.05 | 7030 | 3210 | 0.457 | 0.625 | 0.9103 | 0.71 | 0.798 | 0.650 | 0.836 | 0.367 | 0.64 | None |
| D5-6928 | 84.49 | 43.36 | 1003 | 245 | 0.244 | 0.242 | 0.4425 | 0.31 | 0.520 | 0.550 | 0.464 | 0.508 | 0.51 | None |
| DM-A0XD | 176.37 | 45.13 | 1453 | 306 | 0.211 | 0.285 | 0.9314 | 0.66 | 0.715 | 0.675 | 0.789 | 0.560 | 0.6 | Missense_Mutation |
| DM-A280 | 860.89 | 398.12 | 11938 | 1229 | 0.103 | 0.113 | 0.9301 | 0.4 | 0.540 | 0.750 | 0.542 | 0.457 | 0.47 | None |
| F4-6856 | 424.60 | 174.41 | 4971 | 385 | 0.077 | 0.088 | 0.9244 | 0.6 | 0.777 | 0.800 | 0.752 | 0.616 | 0.63 | Nonsense_Mutation |
| G4-6307 | 410.88 | 215.09 | 5228 | 3369 | 0.644 | 0.656 | 0.9759 | 0.7 | 0.837 | 0.800 | 0.835 | 0.532 | 0.7 | Missense_Mutation |
| G4-6322 | 539.83 | 283.59 | 8410 | 2300 | 0.273 | 0.373 | 0.9237 | 0.88 | 0.752 | 0.800 | 0.825 | 0.760 | 0.72 | None |

*HEMnet tumour purity prediction results are compared to seven genomics based methods
